# Conditioned place avoidance using encapsulated calcium propionate as an appetite suppressant for broiler breeders Conditioned place avoidance and calcium propionate

**DOI:** 10.1101/441220

**Authors:** Aitor Arrazola, Stephanie Torrey

## Abstract

Broiler breeders, the parent stock of meat chicks, are feed-restricte throughout rearing to avoid obesity-related problems in their health and reproductive performance. Broiler breeders often show signs of chronic hunger, lack of satiety and feeding frustration, and the development of alternative feeding strategies has investigated the inclusion of calcium propionate (CaP) as an appetite suppressant. However, the mechanisms involved in the reduction of voluntary feed intake are unknown, but are thought to be due to low palatability, gastrointestinal discomfort, or both. The objective of this experiment was to examine the effect of CaP as an appetite suppressant on the experience of a negative affective state, using a conditioned place preference test. Twenty four broiler breeders were trained to associate the consumption of CaP or a placebo pill with a red or blue place, depending on inherent colour preference. Pullets consumed two pills followed by 20 g feed allotment. The CaP pill contained 160 mg of CaP and the placebo pill had 160 mg of feed. Conditioning lasted for 90 min/pullet/day over 8 consecutive days at 7 and 9 weeks of age, and pullets’ choice was tested in a T-maze twice on two consecutive days at both 8 and 10 weeks of age. Data were analysed using a linear mixed regression model, with pen nested in the model and age as a repeated measure. Pullets were less likely to choose the place conditioned with the consumption of CaP (P<0.05) and the preference of the placebo linearly increased with training sessions (P<0.05). These results suggest that calcium propionate as an appetite suppressant can induce a negative affective state, with the lower feed intake resulting from a conditioned response to the negative effect of calcium propionate rather than to satiety.

## Introduction

Calcium propionate (CaP) is an organic acidifier commonly used in human nutrition as a feed preservative (e.g., bakery [1], in fermented dairy products [e.g., 2]) in animal nutrition as a feed preservative [e.g., 3-6], as a growth promoter [6,7], as an intestinal microbiota enhancer [6,8], and as an appetite suppressant [9–12]. The effect of dietary CaP been researched in poultry because of its role as appetite suppressant. Previous research reported that increasing concentration of CaP (1-3%) linearly decreased feed intake in broiler chickens (i.e., meat chickens) [9], and high inclusion rates of CaP (5-9%) has been proven to lower feed intake in broiler breeders [10–12]. Broiler breeders are the parent stock of broiler chickens and both share the same genetic predisposition for fast growth rate, feed efficiency and high feed intake [13]. Feeding broiler breeders to satiety is associated with obesity-related problems such as low liveability and negative consequences in reproductive performance [14,15]. Therefore, broiler breeder pullets are feed-restricted throughout their production cycle up to 43% of *ad libitum* feed intake for the same broiler body weight, leading to chronic hunger, lack of satiety and frustration [16]. The experience of negative affective states is an ethical concern in animal welfare, and the development of alternative diets has considered the inclusion of CaP as an appetite suppressant to reduce voluntary feed intake and feeding motivation [11,17,18].

Previous experiments have indicated the negative consequences of dietary CaP in humans (e.g., behavioural disturbance [19] and seizures [20]), rats [21], and poultry [12,17]. Daily calcium propionate intake in humans is estimated about 0.1% of food intake [22], although health recommendations allow an inclusion rate up to 0.2% for daily human consumption (e.g., [1,19]). In medical studies, propionic acidemia can result in seizures independent of metabolic acidosis [20] and a controlled trial with children indicated that short-term high inclusion of CaP (at 2% in a bakery product) can lead to irritability, restlessness and sleep disturbance [19]. In the case of poultry, much higher inclusion rates are added in poultry diets compared to human nutrition and previous research noted that chickens avoided consuming the CaP diet if possible [23,24]. For example, Savory *et al.* [23] indicated that the feed intake of broiler breeders (i.e, naïve to CaP) was four times higher with a standard diet compared to a standard diet with CaP. Torrey *et al.* [24] also reported that feed intake of an alternative diet (40% soybean hulls and 3-5% CaP) was low in chickens given free access to both a standard and an alternative diet in their home pen. Afterwards, authors tested the preference for the alternative and the standard diet in a Y maze, and 50% of the non-feed restricted pullets consistently chose the standard diet, with only 6.9% pullets preferentially choosing that alternative diet [24]. Tolkamp *et al.* [12] also observed a high number of oral lesions, and morphological changes in the crop and gastric lesions in broiler breeder fed alternative diets *ad libitum* at 6-9% CaP, while broiler breeders on the standard diet had no lesions. The effect of CaP as an appetite suppressant in poultry has been associated with low palatability, gastrointestinal discomfort/malaise or the combination of both [17,23]. Independent of its palatability, whether CaP causes a negative affective state post-ingestion remains unclear.

The affective states of animals cannot be measured directly, but the study of animals’ choices is an indirect approach to assess their preferences [25]. For example, behavioural tests that looked at the affective states and the preferences of broiler breeders include operant learning/response, state dependent learning and conditioned place preference/avoidance [26–31]. Conditioned place preference/avoid (CPP/CPA) tests are often used to analyse the possible positive (i.e., rewarding) or negative (i.e., punishing) effects of pharmaceutical drugs or dietary additives [32]. During CPP/CPA tests, animals are trained to associate given characteristics of its environment with affective states induced by the treatment, and this environmental cue can become a conditioned stimulus associated with the given affective state [32,33]. Once the condition is created, animals are hypothesized to prefer the environment associated with the relative higher positive affective state when they are tested in a T-maze [32,33]. Indeed, Phillips and Strojan [34] showed that broilers were able to discriminate three pairs of conditioned stimuli for CPP after 7-days conditioning period per each pair. Previous research in broiler breeders has examined the preference between qualitative restriction with CaP and quantitative feed restriction using a CPP [28]. Results in that experiment suggested that chickens failed to learn the CPP due to severe feed restriction [27,30]. However, a lower feed restriction level could facilitate conditioning learning during the training sessions [35]. The objective of this experiment was to examine the causal effect of CaP as an appetite suppressant on a negative affective state by using a conditioned place preference (CPP) test in broiler breeder pullets. Pullets were hypothesized to avoid the place conditioned with the consumption of CaP if the CaP induces a negative affective state.

## Materials and Methods

A total of 24 Ross 308 broiler breeder pullets were donated at one day of age, courtesy of Aviagen (via Horizon Poultry, Hanover, Ontario, Canada). All the procedures used in this experiment were approved by the University of Guelph’s Animal Care Committee (AUP # 3141) and were in accordance with the guidelines outlined by the Canadian Council for Animal Care.

## Housing and management

At the hatchery, chicks were infrared beak treated and vaccinated based on local recommendations, and chicks were raised at the OMAFRA Arkell Poultry Research Station (Guelph, ON, Canada). Chicks were distributed upon arrival to cages (10.3 chicks/m^2^) and were relocated at 22 days of age to six floor pens (4 pullets/pen at 1.1 pullets/m^2^). Floor pens (1.7 m wide × 2.0 m deep × 1.2 m high) had wood shavings as bedding and water was available *ad libitum* from a 5-nipple drinker. Chicks were selected based on target body weight and were individually wing tagged.

Pullets were managed based on breeding company’s guidelines [36]. Room temperature started at 32 °C at 1 day old, and temperature gradually decreased to 29 °C by 2 weeks of age. After transfer to floor pens, room temperature was reduced to 24 °C at 4 weeks of age and to 22 °C at 6 weeks of age. Relative humidity remained around 26% during rearing. Light program was 23D:1D at 100 lux at 1 day old, 12L:12D at 30 lux at 4 days of age until 14 days of age, and 10L:14D at 40 lux until the end of the experiment. Lights came on at 9:00 and pullets were fed simultaneously. Pullets were hand-fed daily using a hanging cylinder feeder with 15 cm/pullet feeder space, and pullets were feed restricted following feed allotment recommendations by breeding company (Table 1;[37]) using standard broiler breeder pullet diet in Starter (0 to 5 weeks of age) and Grower (6 to 10 weeks of age) according to nutrition specifications [38].

**Table 1.**
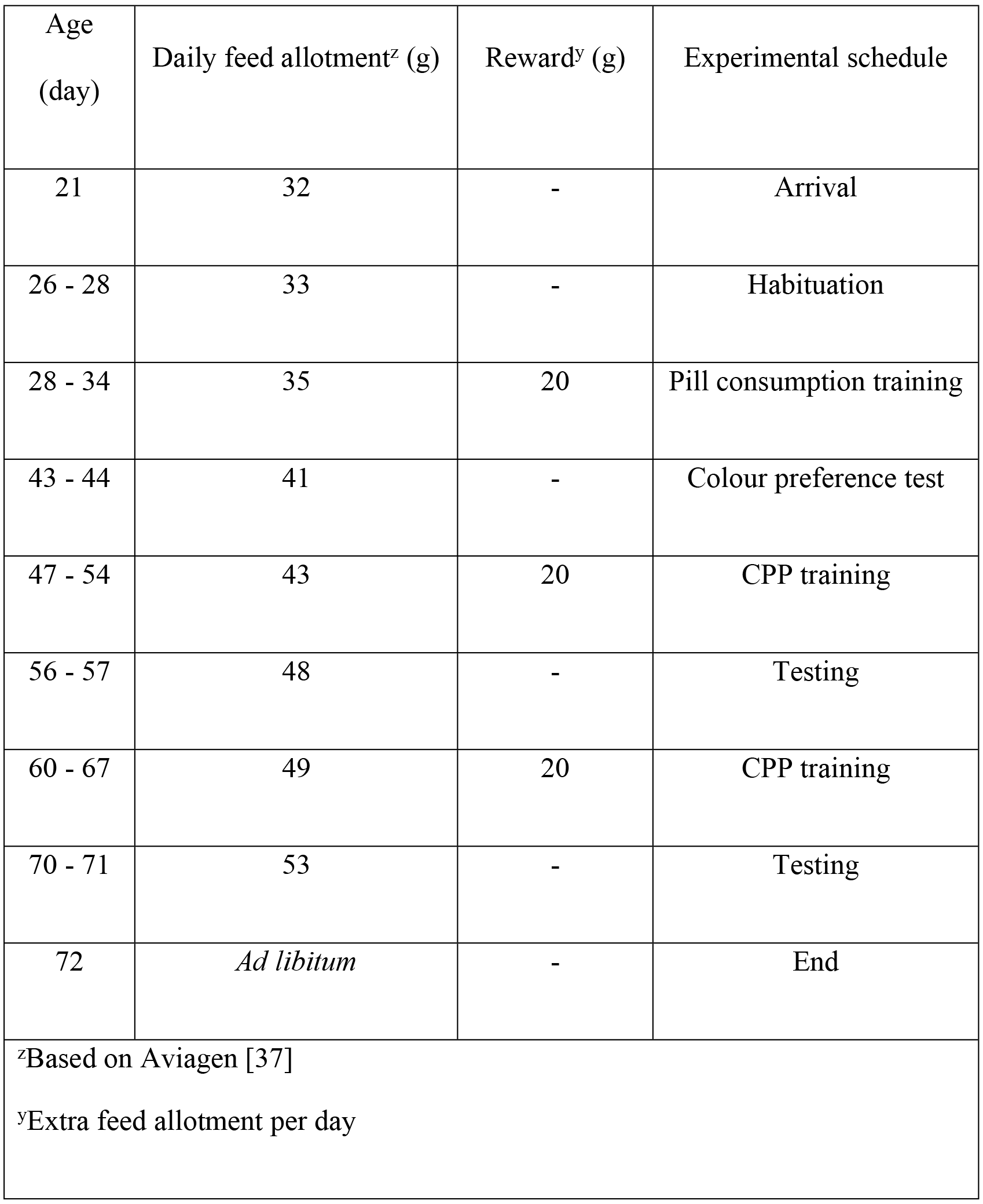
Daily feed allotment and feed reward per pullet depending on age and weekly experimental schedule.

## Experimental design

The experiment lasted from 3 to 10 weeks of age, and pullets were trained to associate the effect of consumption of CaP with a given place accounting for inherent colour preference. Pullets were fed two identical pills (white lock-ring gelatine pill, size 3, CapsulCN): both either CaP or placebo during the CPP training, and pill consumption was rewarded with 20 g of their standard (mash) diet. The CaP pill contained an inclusion of 160 mg of CaP and the placebo pill had 160 mg of their standard diet. In combination with the feed reward, CaP was fed at 1.6% inclusion rate, similar to the inclusion rate of CaP in broiler breeder alternative diets [18]. Placebo pills were used for the voluntarily pill consumption training, the conditioned place preference (CPP) training and for T-maze choice (one pill per place); CaP pills were only used during the CPP training depending on training schedule (Table 2).

There were six pens, and the four pullets per pen were trained simultaneously. Two pullets per pen were trained to associate the CaP pill with the blue place and remaining two pullets per pen were trained to the CaP pill with the red place (Table 2). During the CPP training, pullets alternated pills every two consecutive days for 8 days. Two pullets per pen started the training with the CaP pills, and the other two pullets started with the placebo pills (Table 2). On the two consecutive days with the same pill in two apparatuses (Fig 1), pullets were trained in both sides of the T-maze for the same place colour. For example, if a pullet was trained in the left side the first day with blue walls, the following day, the same pullet was trained on the right side with blue walls, in the other apparatus (Fig 1). The reverse order was used for the other pullets from the same pen (Table 2).

**Table 2.** Experimental design and individual training protocol for the development of conditioned place preference according to pullets’ age. The same design was applied to all pens.

| Age (week) | Pullet id | Colour conditioned with the CaP pill<sup>z</sup> | First pill<sup>y</sup> | First colour<sup>y</sup> | First side<sup>x</sup> | T-maze<sup>x</sup> |
| --- | --- | --- | --- | --- | --- | --- |
| 6 | 1 | Blue | CaP | Blue | Left | 1 |
|  | 2 | Red | CaP | Red | Right | 1 |
|  | 3 | Blue | Placebo | Red | Left | 2 |
|  | 4 | Red | Placebo | Blue | Right | 2 |
| 8 | 1 | Blue | Placebo | Red | Left | 2 |
|  | 2 | Red | Placebo | Blue | Right | 2 |
|  | 3 | Blue | CaP | Red | Right | 1 |
|  | 4 | Red | CaP | Blue | Left | 1 |
| <sup>z</sup>Colour was selected based on the initial preference before training started<br><sup>y</sup>Pullets remained on this pill/colour for two consecutive days<br><sup>x</sup>Alternated every other day |  |  |  |  |  |  |

**Fig 1.**
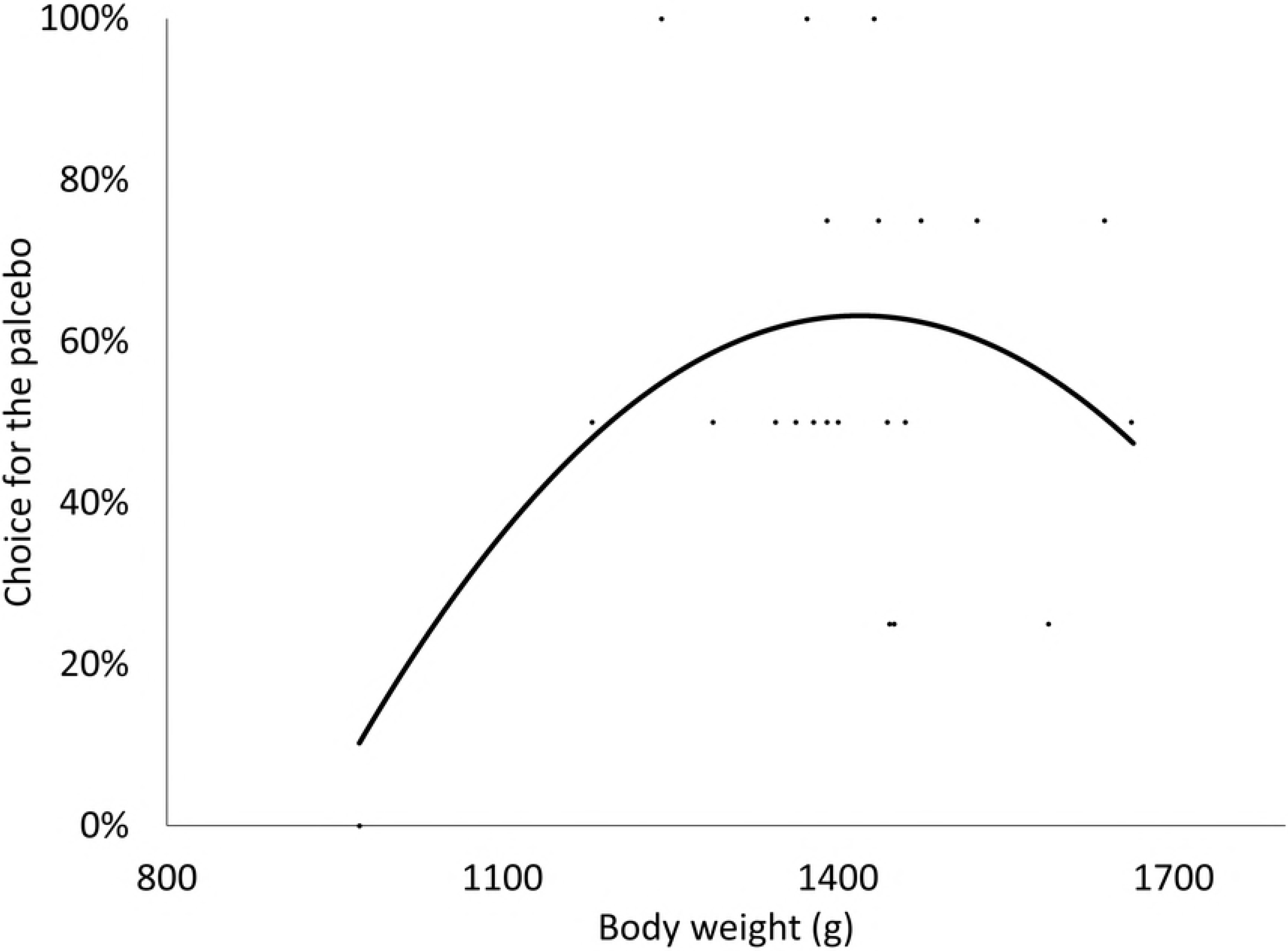
Diagram of the T-maze used for the inherent colour preference, for the conditioned place preference and the choice test. Pullets were trained in each arm for conditioned place preference, visually isolated from each other during the CPP training. Two mirror-image apparatuses were used, with colour location differing between arms (left versus right side). Diagram is drawn to scale (1:8.5).

The choice and preference for the placebo was tested in the T-maze (Fig 1). Fig 1 represents the apparatus used for the CPP and the T-maze of the choice test. Two apparatuses were used in the experiment and each one was located inside a pen similar to pullet’s home pen within the same room. Both apparatuses had the same dimensions (145 cm wide × 132.1 cm deep × 48.3 cm height) but differed in the location of the blue and red arms (right versus left side). Each pullet per pen was trained to associate the effect of the consumption of CaP with one of the colours, controlling for side. On testing days, one placebo pill was offered in each arm of the T-maze, but pullets could only consume one of them in each testing event. The operant response in this experiment was the consecutive consumption of two pills and pullets were positively rewarded afterwards; however, pill consumption was not rewarded on testing days due to only one pill being accessible in each testing event.

## Methodology

This research was divided in four phases: 1) habituation, 2) individual colour preference, 3) CPP training, and 4) choice test. Habituation to the T-maze, training for pill consumption, place preference test, conditioned place training, and testing started after all pullets consumed their daily feed allowance. The preference and choice tests started one hour after feeding, and the conditioned place training started one hour after feeding for the first pen to seven hours and half for the last pen. The average body weight (± SD) was 366.7 ± 34.8 g, 1075.0 ± 114.9 g and 1408.5 ± 144.9 g at 3, 8 and 10 weeks of age, correspondently.

## Habituation

Pullets were trained for voluntary and consecutive consumption of two placebo pills during the first week of habituation. On the first day, pills are manually fed and consumption of two pills was rewarded with 20 g of the standard diet. Afterwards, two placebo pills were simultaneously offered per pullet per day for seven days, until pullets voluntarily consumed both pills when offered. Pullets were rewarded with 20 g of feed if pills were manually fed or voluntarily consumed.

Pullets were placed inside the testing pens where the T-maze was located, and pullets could explore the T-maze for 2 hours to habituate them to training and testing conditions. Habituation to the T-maze started at 4 weeks of age and lasted for 10 days. Initially, the four pullets per pen were grouped within the same testing pen and the T-maze. After the second exposure, habituation was done in pairs. At the end of the first week, pullets were individually placed in the testing pen for 2 hours to habituate them to isolation. Once pullets were individually isolated, two placebo pills were provided, and pill consumption was rewarded with 20 g of the standard diet.

## Inherent bias test (Pre-CPP)

After habituation and training, each pullet was tested on two consecutive days and twice per day to assess individual preference for side and colour at 34 days of age. During the preference tests, pullets saw both T-maze arms (coloured) for the first time. The order of pullets in this test was randomly chosen and pullets were retested after all pullets were tested once on the same day (no one was tested consecutively). Another random pattern was generated for the following testing day. Pullets were placed into the starting box for 30 sec, and then the door to the maze was opened. An observer registered pullets’ inherent choice, defined as the whole body of the pullet inside one of the sides of the T-maze for more than 5 seconds. Pullets had 5 minutes to choose either left or right, otherwise they were registered as having no choice (assuming no colour preference). Colour and side preference was set for pullets that chose one colour or side at least three times out of four trials (≥ 75%). All pullets were tested on the same day in both apparatuses and the order of the starting apparatus was reversed for the second day of testing.

## CPP training

The CPP training lasted for eight consecutive days per training session. All pens were trained daily and the four pullets per pen were training simultaneously, one in each conditioned place. Two pullets per pen were trained to associate the placebo pills with the blue walls and the CaP pills with the red walls, and vice versa for the other two. Pullets that had an inherent bias for a colour before the CPP training started received the placebo pills in the opposite colour to their preference (except for two pullets for which three pullets from the same pen showed preference for the same colour). The CPP training sessions were repeated twice at 7 and 9 weeks of age, and pullets were tested for preference choice for the placebo place at the end of each training session.

Each pullet was individually allocated in the conditioned place for 90 min. Pullets were trained to consume two pills in 30 seconds and were rewarded with 20 g of the standard diet after the consumption of the second pill. Pullets had *ad libitum* access to nipple drinkers in the apparatus during the conditioned phase and were visually isolated from the other conditioned place.

## Choice test

Pullets’ choice was tested individually on two consecutive days using the same protocol for the inherent bias test (i.e., same testing order and methodology) at 8 and 10 weeks of age one day after the last day of each training session. During the choice test, one placebo pill was left in each T-maze arm and each pill was set in the middle of the conditioned place (equidistant and visible from the fork). An observer recorded the place from which pullets consumed the pill. Pill consumption was not feed-rewarded and pullets were returned to their home pen after consuming one pill. Data were collected according to pullet’s choice and preference for the placebo pills. Choice for the placebo was defined as the percentage of times that pullets consumed the pill from the placebo place. Preference for the placebo was defined as the percentage of pullets that chose the placebo place consistently (≥75%) divided by the total number of pullets tested.

Individual body weight was recorded on the last testing day at 8 and 10 weeks of age. After all pullets were tested at 10 weeks of age, pullets were individually presented with four pills (two placebo pills and two CaP pills) to examine whether pill consumption would stop after the consumption of the second pill and whether pullets could differentiate between both pills. Both pills were visually identical and had the same weight, but pills might have differed for pullets’ perception. In addition, pullets were fed *ad libitum* afterwards in their home pen and remaining placebo pills were offered in the feeders.

## Statistical analysis

The choice for the CPP was statistically analysed using a generalized mixed linear model and pen was nested in the model as the independent research unit. The statistical analysis was performed using SAS Ver. 9.4 (SAS Institute, Cary, NC) with a glimmix procedure and the degree of significance was set for p-values lower than 0.05.

Age was included as a fixed factor in the model for the choice and preference for the placebo. Initial observations (i.e, 3 weeks of age) for the choice and the preference for the placebo were considered as independent observation rather than a covariate to achieve desirable degrees of freedom as well as for orthogonal regression analysis for age. The effect of the last training day was included as a covariate. Similarly, the effect of body weight was included as a covariate within age due to collinearity. The assumptions to the analysis of variance were assessed using scatterplots of studentized residuals, linear predictor for linearity, and a Shapiro-Wilk test for normality. Outliers were defined as observations with absolute studentized residuals higher than 3.4. Individual differences, pen, and pen location within the room were included in the covariance structure as random effects. Age was fit into a repeated structure for placebo choice with pen as the subject with age as the group. Orthogonal regressions analysed the effect of age into linear and quadratic response. Significance differences between multiple mean comparisons were corrected using Tukey-test adjustment.

## Results

One pullet was excluded from the dataset due to unsuccessful training to voluntarily pill consumption. Pullets were 62.9% and 32.8% heavier than the performance objectives for broiler breeders at 8 and 10 weeks of age, respectively [37] due to feed rewards. Pullets were trained to associate the effect of CaP with the colour they inherently preferred. However, all pullets from one pen preferred the red colour and for two of these pullets, the placebo remained in the colour they preferred due to consistency with the training protocols. Hence, the initial choice for the placebo was higher than 0% (Fig 2). Data is presented using estimated mean values followed by the standard error of the mean.

**Fig 2.**
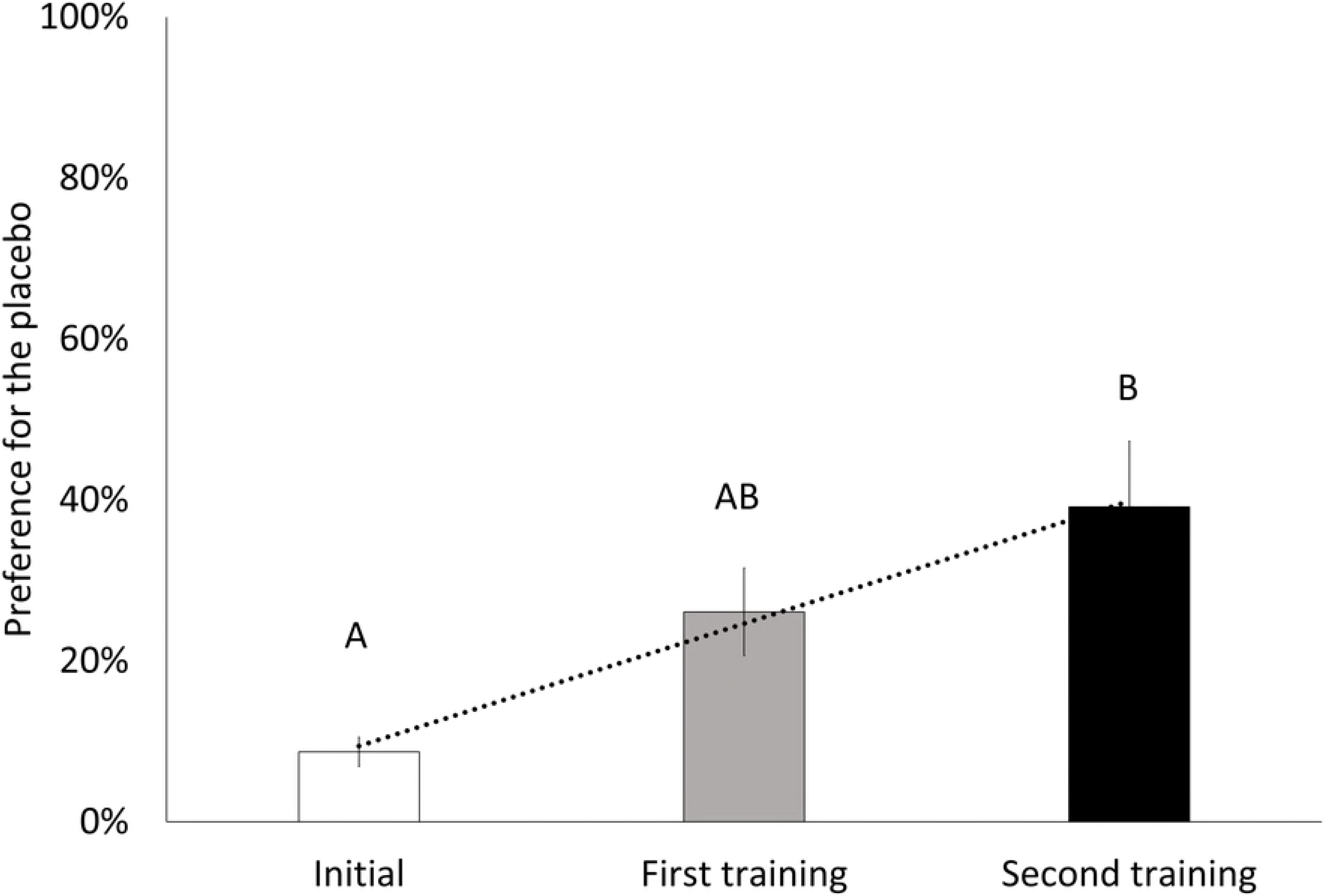
The effect of conditioned place preference training in broiler breeder pullets over time on the choice (A) and preference (B) for the placebo pill compared to the calcium propionate pill (means ± SE) The choice of the placebo indicates the proportion of times that the placebo place was chosen (A). The preference for the placebo represents the proportion of hens that consistently chose the placebo place divided by the total number of hens (B). Pullets were tested on day 42 for initial choice, on day 56 after the first training session and on day 70 at the end of the second training session. Repetitive training sessions linearly increased the choice (F_1,18_=60.90, P<0.001) and preference (F_1,18_=38.27, P<0.001) for the place conditioned with placebo pill.

## Individual colour and side preference

Pullets showed a significant preference (choosing ≥75% of times) for colour and side at 6 weeks of age. Half of the pullets (12 out of 24) preferred the red place over the blue place (t=4.49, P<0.001) and only one pullet out of 24 preferred the blue place over red. Three pullets showed lateralization during the choice preference test (t=4.89, P<0.0001); two pullets preferred the right side and the one preferred the left side. Moreover, four pullets out of 24 chose no place during the initial preference test.

## Choice test

All pullets chose a place at each time they were tested at 8 and 10 weeks of age. During the choice test, 17.4% (4 pullets out of 23) and 13.1% (3 pullets out of 23) of pullets showed consistent lateralization (choosing either left or right the four times) at 8 and 10 weeks of age, respectively; and only one pullet was consistent toward the left side at both ages (trend not observed during the initial side preference). The choice for the placebo was influenced by body weight at 70 days of age (F_1;22_=5.32, P=0.03) and body weight had a quadratic effect on the choice of the placebo at 10 weeks of age (Fig 3). The pullets that preferred the placebo place ranged from 1242 g to 1432 g at 10 weeks of age whereas pullets outside this threshold showed no preference. Between the place conditioned with the consumption of the placebo or the CaP pill, pullets only showed preference for the placebo place. Individual variability did not affect the choice for the placebo (Z=1.00, P=0.16). As well, the last training day before the T-maze choice did not influence the choice (F_1;44_=0.74, P=0.40) nor the preference (F_1;44_=2.20, P=0.16) for the placebo during the T-maze tests.

**Fig 3.**
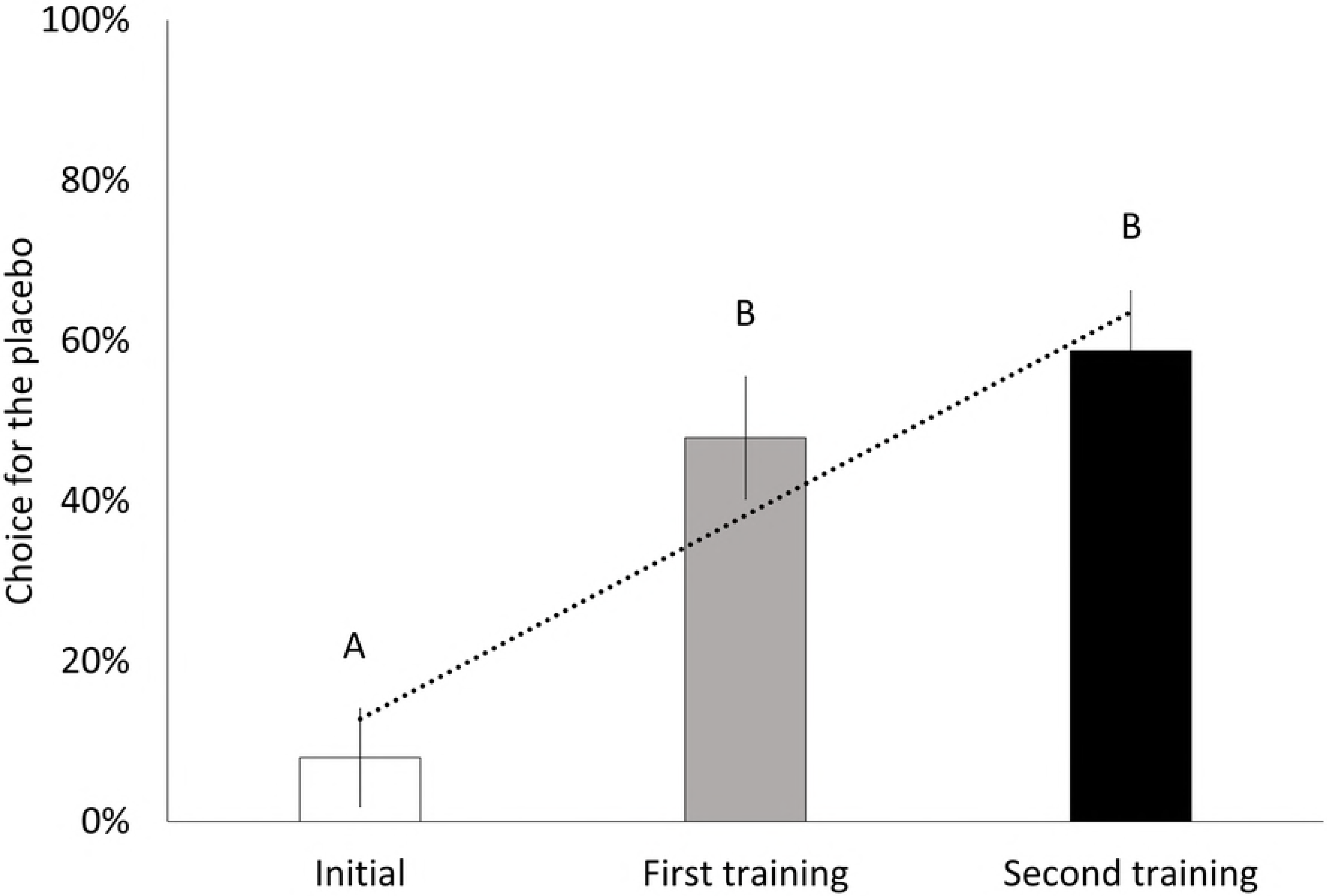
The effect of body weight on the choice for the place conditioned with the placebo pills at 70 days of age. Body weight had a quadratic effect on the choice for the placebo at 10 weeks of age (F_1,22_=5.32, P=0.03).

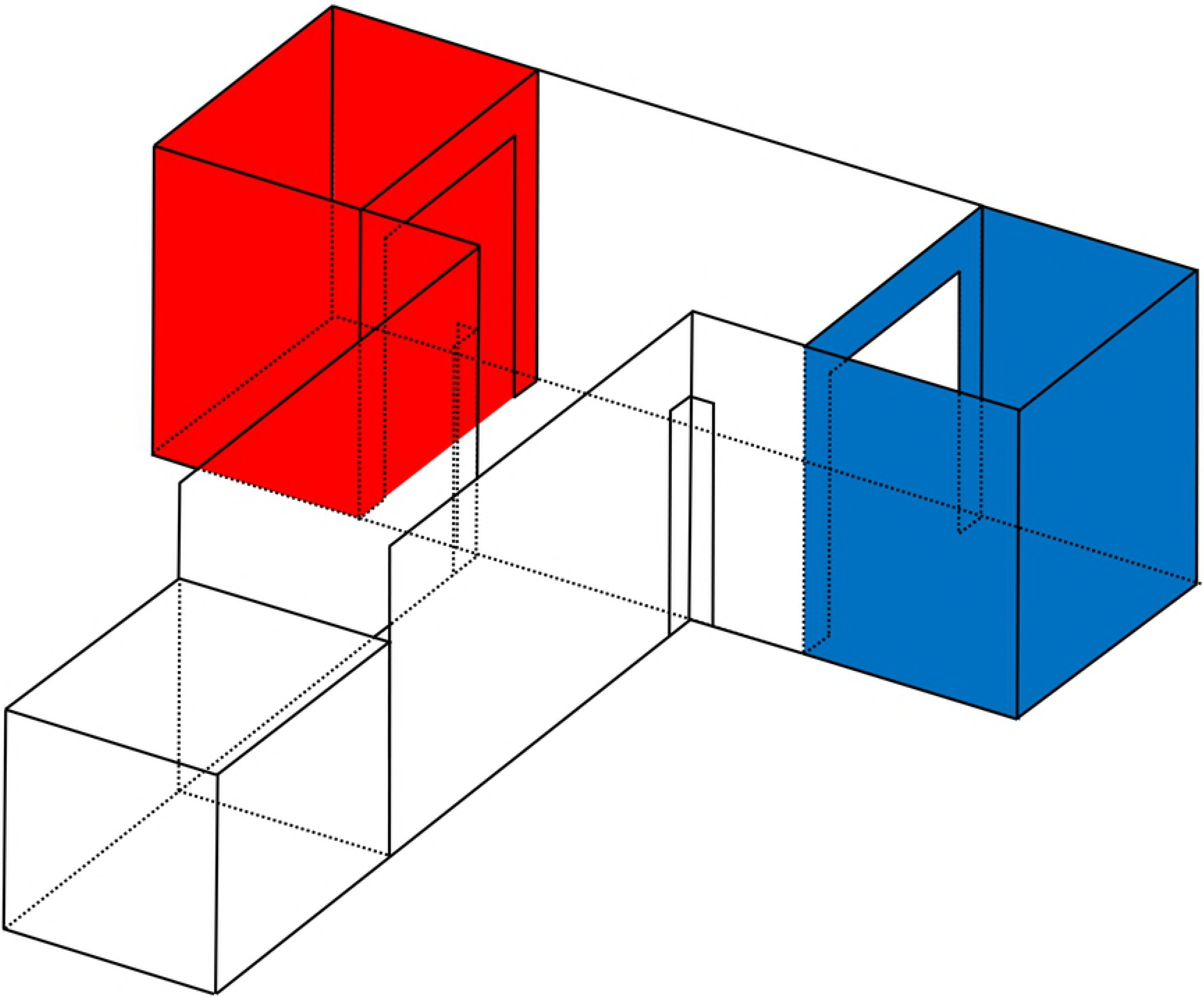

Fig 2 illustrates the effect of training on the choice (F_2,18_=36.07; P<0.0001) and the preference (F_2,18_=4.19; P=0.03) of pullets toward the placebo pill. Repetitive training sessions linearly increased the probability for the pullets to choose the place conditioned with the placebo pill (F_1,18_=60.90, P<0.001). The probability for the pullets to choose the placebo increased from 8.0 ± 6.1% at 6 weeks of age (which was artificially manipulated to be as close to 0% as possible) to 47.9 ± 7.6% at 8 weeks of age (t_18_=4.68, P=0.0005), and to 58.7 ± 7.5% at 10 weeks of age (t_18_=6.06, P<0.001). Pullets preferred the place conditioned with the placebo pills after the first training session at 8 weeks of age (25.0 ± 8.6% [6/23]; t_18_=2.91, P=0.01) and at 10 weeks of age (38.9 ± 8.6% [9/23]; t_18_=4.53, P<0.001). The preference for the placebo linearly increased over time (F_1,18_=38.27, P<0.001), and the preference for the placebo increased by 30.56 ± 10.57% from 6 to 10 weeks of age after two training sessions (t_18_=2.89, P=0.03). Moreover, all pullets indiscriminately consumed the four pills in less than 30 seconds at the end of the choice test at week 10 and pullets still ate the placebo pills after *ad libitum* feeding.

## Discussion

The present study was designed to determine the causal effect of the CaP as an appetite suppressant on a negative affective state. A CPP test was conducted using encapsulated CaP at a similar inclusion rate as in alternative diets for broiler breeders [18]. Pullets were expected to avoid the place conditioned with the consumption of CaP if CaP induces a negative affective state, and our results indicate that pullets avoided the place conditioned with the consumption of CaP and developed preference for the placebo place over time. This study demonstrates the causal effect of CaP on inducing a negative affective state at an inclusion rate proven to decrease voluntary feed intake in poultry. Preference was only shown toward the placebo after CPP training and 9 out of 23 pullets that showed a preference chose the placebo place at the end of the experiment (as illustrated in Fig 3). Savory *et al.* [23] examined the dietary preference for CaP at 5% inclusion in broiler breeder pullets, and the feed intake among diets differed based on pullets’ previous experience with CaP. In their research, pullets reared on the standard diet consumed more of the standard diet (81% standard diet vs 19% CaP diet) compared to pullets already on the CaP diet (65% standard diet vs 35% CaP diet). The authors concluded that the preference for the diet at 5% CaP was low, although pullets reared at a fixed CaP inclusion rate might have been habituated [23]. Therefore, the effect of CaP related to a post-ingestive and dose-dependent mechanism, not acidity, to which pullets can habituate [23]. Although the decrease in feed intake may be due to a lower palatability of the diet with CaP [23,24,39], CaP was encapsulated in our research to avoid the confound between palatability and a post-ingestive mechanism. Similar results on the aversiveness of CaP were reported in rats in Ossenkopp *et al.* [21]. The authors infused CaP (at 0.05%) to avoid the confound of palatability, and rats spent less time in the place conditioned with the CaP, and showed more escape attempts and hyperactivity, indicators of aversiveness, during conditioning training. In agreement with our results, CaP can induce a negative affective state regardless of its palatability, although CaP may have an additional effect at reducing voluntary feed intake due to low palatability [21]. Therefore, the effect of CaP as an appetite suppressant can relate to a conditioned response to the causation of a negative affective state instead of satiety.

Our results illustrate that training had a linear effect on the consistency of instantaneous choices, suggesting that feed-restricted pullets may require more and longer training for the condition to happen. Dixon *et al.* [30] concluded that broiler breeders under commercial feed restriction were unsuccessful in learning CPP or CPA tests, and chronic feed restriction can limit cognitive potential. Previous research indicated that chronic stressful conditions such as feed restriction could limit associative learning [27,40]. Certainly, broiler breeders display behavioural and physiological signs of chronic feed restriction [16], and commercial feed restriction can be distressful for broiler breeders [15]. Buckley and colleagues [40] examined whether broiler breeders can learn a discrimination feed intake test *(i.e.,* pullets were trained to associate the arm colour with the amount of feed reward) depending on feed restriction level (commercial feed restriction, 40% and 80% of *ad libitum* feed intake). Buckley *et al.* [40] indicated that the latency to perform the choice test decreased as the feed restriction level increased, and a lower proportion of pullets learned the CPP tests on the highest feed restriction level (i.e., commercial feed restriction level). The deficient performance of the pullets under commercial feed restriction can be explained by their inability to learn during the CPP training or arousal during the choice test due to light body weight, high feeding motivation and chronic stress due to severe feed restriction. Additionally, Buckley *et al.* [28] examined the preference of broiler breeders under commercial feed restriction between qualitative restriction (at 3-9% CaP) and quantitative feed restriction, but the authors did not observe differences between both treatments probably due to lack of preference or inability to learn under such severe feed restriction [28]. In agreement with our results, Buckley *et al.* [28] and Dixon *et al.* [30] also suggested that broiler breeders might require numerous and long training sessions to facilitate associative learning and to control for high feeding motivation under commercial feed restriction. For this reason, the feed restriction level was lowered in our experiment and pullets in this research exceeded target body weight because of multiple feed rewards during training for pill consumption and for the conditioned place learning.

Body weight varied widely in our research at 10 weeks of age, and lighter and heavier pullets were less likely to choose the placebo place compared to pullets of average body weight. Previous research also reported that lighter broiler breeders performed poorly in a learning task [28,30,40]. Results in Buckley *et al.* [40] pinpoint that (light) pullets under commercial feed restriction performed poorly in a learning task compared to (heavy) pullets reared on a lower feed restriction level. In our experiment, pullets were more likely to choose the place conditioned with the placebo at a body weight between 1200 g and 1450 g at 10 weeks of age, which is approximately 100-350 g heavier than target body weight. Feed restriction was the same among pen mates in our experiment, but pullets were group housed and fed via a communal feeder. Variation in body weight among pen mates may reflect previous experience in their home pen [41] as well as current feeding motivation [39], resulting in a high arousal during behavioural tests [40–42]. Lindholm *et al.* [41] reported that pullets at light body weights were more active in a tonic immobility test. In Buckley and colleagues [40], pullets under commercial feed restriction had shorter latencies than pullets fed at 80% of *ad libitum* feed intake in a T-maze choice test. Previous research with broiler breeders also noted that fearfulness linearly decreased as feed restriction increased [14,15] and during off-feed days under non-daily feed restriction [43]. For this reason, light broiler breeders under feed restriction conditions display high arousal as suggested by short latency and low fearfulness in behavioural tests [40–42], probably being less consistent in short-term choices due to previous experiences and current feeding motivation. On the other side, heavy pullets were not as likely to choose the placebo place as average weight pullets in our experiment, and one of the heaviest pullets was the only one to show consistent lateralization at 8 and 10 weeks of age. Buckley *et al.* [40] noted that broiler breeders displayed lateralization in their decision-making behaviour during a choice test depending on the feed restriction level. All 12 pullets raised under commercial feed restriction showed lateralization compared to only 6 of 12 pullets at 80% of *ad libitum* feed intake. Buckley *et al.* [40] indicated that lateralization can relate to hunger stress, with hungrier pullets being more resistant to change behaviour (lower behavioural plasticity). In Arrazola [39], we also observed that heavy pullets were more highly feed motivated compared to lighter pen mates at the same age and feed restriction. For this reason, heavy (and dominant) pullets may be more highly motivated compared to pullets of average body weight, contributing to arousal and inconsistency during a choice test. However, the role of feeding motivation, body weight and individual variation in decision-making behaviour is unknown, despite its potential to improve body weight uniformity under commercial conditions in broiler breeder houses. Anecdotally, pullets showed short latencies during the choice test, and consumed both pills rapidly during the CPP training. The choice test in our research was not performed under extinction conditions, although the T-maze choice was not rewarded. Thus, visual cues (i.e., the pill in our research) can be an external factor that may have triggered arousal in pullet’s choice, especially in light and heavy pullets at 10 weeks of age. Elevated feeding motivation (either in light and heavy pullets) can interfere in decision making behaviour and compromise the performance during the choice test of broiler breeders [27,30,40,41], which may explain the effect of body weight in our research. For this reason, previous and current experiences should be considered in the interpretation of results about animals’ preference because high feeding motivation and arousal state can be translated into inconsistency or lack of preference in their choice. The effect of body weight can be alternatively explained by pullets’ inability to learn the CPP due to chronic stress (35). However, behaviour was not recorded during the conditioning phase or during the choice test in this experiment to assess whether this hypothesis holds true for lighter and heavier pullets. But, further research looking at whether the poor performance in learning task relates to learning inability or arousal by feeding motivation is necessary to properly understand results from preference tests in feed-restricted animals.

T-mazes are used in behavioural tests to examine the animals’ preference between two options [25]. However, a choice may represent something to be avoided rather than something to be desired, such as in our preference test. In this case, both choices were feed restriction with or without the inclusion of CaP. Therefore, these results indicate that the inclusion of CaP (i.e., qualitative restriction) is avoided, instead of commercial feed restriction (i.e., quantitative feed restriction) being preferred. Additionally, results from preference tests rely on previous experiences and training protocol [25]. Dixon *et al.* [30] highlighted that the side of the T-maze was a main driver in the preference of feed-restricted broiler breeders and pullets preferred the place they were not previously housed. The side of the T-maze was considered in the experimental design, and the effect of the last training day on the choice or the preference during the choice test was not significant in our experiment. Inherent colour preference was also considered in our experimental design because colour preference in avian species has been previously reported by Ham and Osorio [44]. Broiler breeders showed an inherent colour preference toward the red colour over blue at six weeks of age as previously described Taylor *et al.* [45] and Fischer *et al.* [46]. However, this preference switched to the place conditioned with the consumption of the placebo pill after two training sessions regardless of inherent colour preference. Therefore, lowering feed restriction (compared to commercial feed restriction) during training and controlling for inherent colour preference may have facilitated the avoidance response toward the place conditioned with the CaP during the choice test.

These results highlight the negative welfare consequences of CaP as an appetite suppressant in alternative diets for broiler breeders, but also as a feed additive in poultry diets. CaP is often used as a growth promoter and feed preservative [6] at concentrations from 0.25% to 0.6% CaP in standard diets [3–5]. Paul and colleagues [4] reported a lower feed intake at 0.5% CaP in one-month-old broilers that resulted in an improvement in the feed efficiency, and similar results were reported by Bonos *et al.* [5] with a standard diet at 0.1% CaP. However, the effect of CaP on feed and intestinal microbial count (Coliforms, *E. coli, Clostridium spp, and Aspergillus spp.)* is not as evident at such inclusion rates [4]. The fungicidal properties of CaP have been previously indicated at 1% inclusion rate, a concentration at which CaP can inhibit the germination and proliferation of *Aspergillus sp.* [47]. For this reason, a similar inclusion rate of CaP to the one used in the previous experiment might be required to achieve the antimicrobial properties of CaP in standard poultry diets. However, our results suggest that the effect of CaP as an appetite suppressant can induce a negative affective state at 1.6% inclusion rate, making its use in poultry, animal and human nutrition controversial from a welfare perspective.

## Conclusion

Pullets were more likely to avoid the place conditioned with the consumption of calcium propionate and pullets developed an increasing preference for the placebo place over time. These results indicate that the inclusion of calcium propionate as an appetite suppressant (at 1.6%) can cause a negative affective state, and this aversive effect can underlie the reduction in feed intake in diets supplemented with calcium propionate. As well, the effect of feeding motivation on arousal and on learning ability on the performance of broiler breeders in learning task or choice test should be considered in the interpretation of animals’ preferences.

## Acknowledgements

Financial support for this project came from the Canadian Poultry Research Council (PWB080), the Poultry Industry Council (Project #325), the Canadian Hatching Eggs Producers, the Natural Science and Engineering Council of Canada (CRDPJ-470617-14), Ontario Ministry of Agriculture, Food and Rural Affairs (Project #030093). The authors thank the Arkell Poultry Research Station personnel for their diligent care of the birds used in this trial and the Animal Behaviour and Welfare group at the University of Guelph for their advice and help.

